# SpatialClaw: A Memory-Augmented Autonomous Ecosystem for Spatial Omics Analysis

**DOI:** 10.64898/2026.05.21.723451

**Authors:** Gaoyuan Du, Ou Lan, Xinrong Wei, Yingfu Wu, Gaotian Meng, Jialun Wu, Zhanhuai Li, Xingyi Li, Xuequn Shang

## Abstract

While the expansion of spatial omics has revolutionized our ability to dissect tissue architecture, the accumulation of incompatible computational methods has heavily fragmented end-to-end analysis, rendering complex workflows irreproducible. Generic conversational agents lack the domain-specific precision necessary to navigate the intricate biological pipelines. To overcome this, we present SpatialClaw, a memory-augmented autonomous ecosystem to unify spatial omics analysis under a single natural-language interaction. SpatialClaw integrates 30 specialized skills, spanning raw data preprocessing, spatial domain identification, deconvolution, spatially variable gene detection, cell–cell communication analysis, multi-sample and cross-modality integration. Distinct from existing agents, SpatialClaw introduces a graph-based persistent memory architecture that stores dataset metadata, analysis lineage, biological insights, and user preferences as versioned nodes and edges across three hierarchical layers (Session, Episodic, and Semantic), governed by a deterministic promotion policy. A Memory-Augmented Reasoning (MAR) Operator bridges the memory store and the main agent, synthesizing retrieved experiences into task-specific guidance for each query. In rigorous benchmarking spanning three memory-sensitive scenarios across 10 spatialomics skills, SpatialClaw outperforms both a standard large language model and the memory-only configuration. Furthermore, we demonstrate its robust biological utility by dissecting the complex tumor microenvironment of a 15-section human triple-negative breast cancer cohort. In merely three conversational turns and with zero direct scripting, SpatialClaw executes a comprehensive end-to-end workflow, yielding standardized output bundles. Ultimately, by synergizing comprehensive analytical tools with structured persistent memory, SpatialClaw elevates spatial omics from disjointed computational stitching to a fully traceable, reproducible, and self-improving discovery ecosystem. SpatialClaw is ready to use at https://github.com/ShangBioLab/SpatialClaw.

## 1 Introduction

The advent of high-throughput sequencing has progressively expanded omics analysis from the measurement of bulk gene expression to the interrogation of cell-state heterogeneity and, most recently, to the spatial organization of gene expression within tissues [1, 2]. Bulk RNA sequencing reports population-averaged expression and obscures cellular heterogeneity, while single-cell RNA sequencing resolves cell types and states but discards the tissue context in which those cells operate [3]. Spatial omics fills this gap by acquiring expression profiles together with the spatial coordinates of each measurement, yielding a joint readout of expression and tissue architecture [4, 1]. Spatial data have proven essential for studying tumor microenvironments, developmental gradients, immune infiltration, and the cytoarchitecture of the nervous system, and the rapid maturation of platforms such as 10x Genomics Visium and Xenium, MERFISH, Slide-seq, and Stereo-seq has steadily increased the scale and resolution at which these questions can be addressed [2].

The growth of the field has been accompanied by an equally rapid proliferation of computational methods spanning the full analytical chain [3, 5, 6]. At the preprocessing layer, tools for quality control, normalization, denoising, batch correction, and spatial smoothing are continually introduced. At the tissue level, methods for spatial-domain identification, spatially variable gene detection, region segmentation, and spatial autocorrelation analysis dissect organizational patterns within slides [7, 8, 9, 10]. At the cellular level, deconvolution, cell-state inference, and cell–cell communication analysis enrich the interpretation of spot-resolved data [11], while joint analyses with single-cell omics, histology, and other modalities push the field toward integrative readouts. Yet these tools are dispersed across heterogeneous platforms, programming languages, and input–output conventions, with no unified analytical framework or standard interface; researchers must repeatedly switch tools, reformat data, and manually thread successive analyses, which raises the technical barrier and limits reproducibility, cross-project portability, and method-level integration.

Large language models—GPT-4, Claude, and their peers—have recently delivered substantial gains in compositional reasoning, planning, and tool use, propelling the evolution of LLM agents from question-answering systems into autonomous task-execution frameworks. Through general-purpose scaffolds such as LangChain and AutoGPT, agents can now decompose complex scientific tasks into multi-step plans, invoke external tools, and manage intermediate state, with particular promise for multi-omics and spatial-omics analysis. Domain-specialized systems have begun to operationalize this potential: OmicClaw [12] integrates more than two hundred analysis functions through a unified multiomics ecosystem (OmicVerse) and a registry-driven engine; ChatSpatial [13] enforces schema-driven orchestration that constrains LLM tool calls to pre-validated interfaces and harmonizes parameter spaces across Python and R; and SpatialAgent [14] organizes experimental design, analysis, and multimodal integration into a memory–plan–execute loop. Despite this progress, existing bioinformatics agents face two systematic limitations in real research scenarios. (i) *Tool coverage remains incomplete*: most systems concentrate on standard pipelines or single-omics tasks, and spatial-omics-specific operations together with multi-omics integration—multi-sample integration, spatial deconvolution, cross-modality fusion, and statistical inference—remain under-represented; tool extension typically depends on manual registration or pre-defined schemas and lags new method publications. (ii) *Persistent memory is limited* : most agents rely on the short-term context window and lack durable storage of dataset state, analysis parameters, historical results, and reasoning trajectories [15, 16, 17]. Even systems that incorporate template memory or task context struggle to maintain continuity across sessions or projects [18, 19, 20], forcing repeated data uploads and reconfiguration that erode the efficiency and reproducibility of multi-step, multi-stage analyses.

To address these limitations, we present **SpatialClaw**, an end-to-end LLM-agent autonomous ecosystem for spatial omics and adjacent multi-omics analysis. Our contributions are twofold. First, we deliver a comprehensive integration of spatial-omics methods. Built on a unified registry, the autonomous ecosystem encapsulates common spatial-analysis tasks—preprocessing, spatial-domain identification, deconvolution, spatially variable gene detection, multi-sample integration, and others—as reusable modules and incorporates established third-party implementations (Tangram, Stereoscope, SpaGCN, STAGATE, GraphST, BANKSY, SpatialGlue, Harmony, Scanorama, DESeq2, CellPhoneDB, and CellChat, among others) to provide end-to-end coverage. The current release ships 30 registered skills, of which 29 are dedicated to spatial omics, exposed through standardized inputs and outputs and dispatched by a single conversational layer that maps natural-language requests onto reproducible multi-step pipelines. Second, we introduce a graph-based persistent memory architecture. Inspired by distinctions between episodic and semantic memory in cognitive science [21] and recent advances in structured memory for LLM agents [22, 23, 24], we model sessions, datasets, user preferences, and analytical conclusions as versioned nodes and edges in a graph, supporting lineage tracing and semantic retrieval, and we expose the subsystem through a high-level API that can also be deployed as a stand-alone memory service. By design, the store retains only metadata rather than raw expression matrices, balancing reproducibility against data privacy, and enables checkpoint-style resumption and result reuse across sessions. Together, these two design choices elevate spatial-omics analysis from fragmented tool stitching to a traceable, reproducible, and extensible end-to-end intelligent autonomous ecosystem, addressing the gaps in tool coverage and persistent memory that have constrained existing systems.

## 2 Results

### 2.1 Overview of the SpatialClaw autonomous ecosystem

The proliferation of spatial-omics methods has produced a fragmented computational landscape in which researchers must manually chain tools written in different languages and frameworks, with no unified execution path or persistent record of prior work. SpatialClaw addresses this by organizing the entire analytical stack into three tightly coupled layers: a single LLM agent as the conversational controller, a curated registry of modular spatial-analysis skills as the execution backend, and a graph-based persistent memory system maintaining cross-session state (Fig. 1).

**Figure 1:**
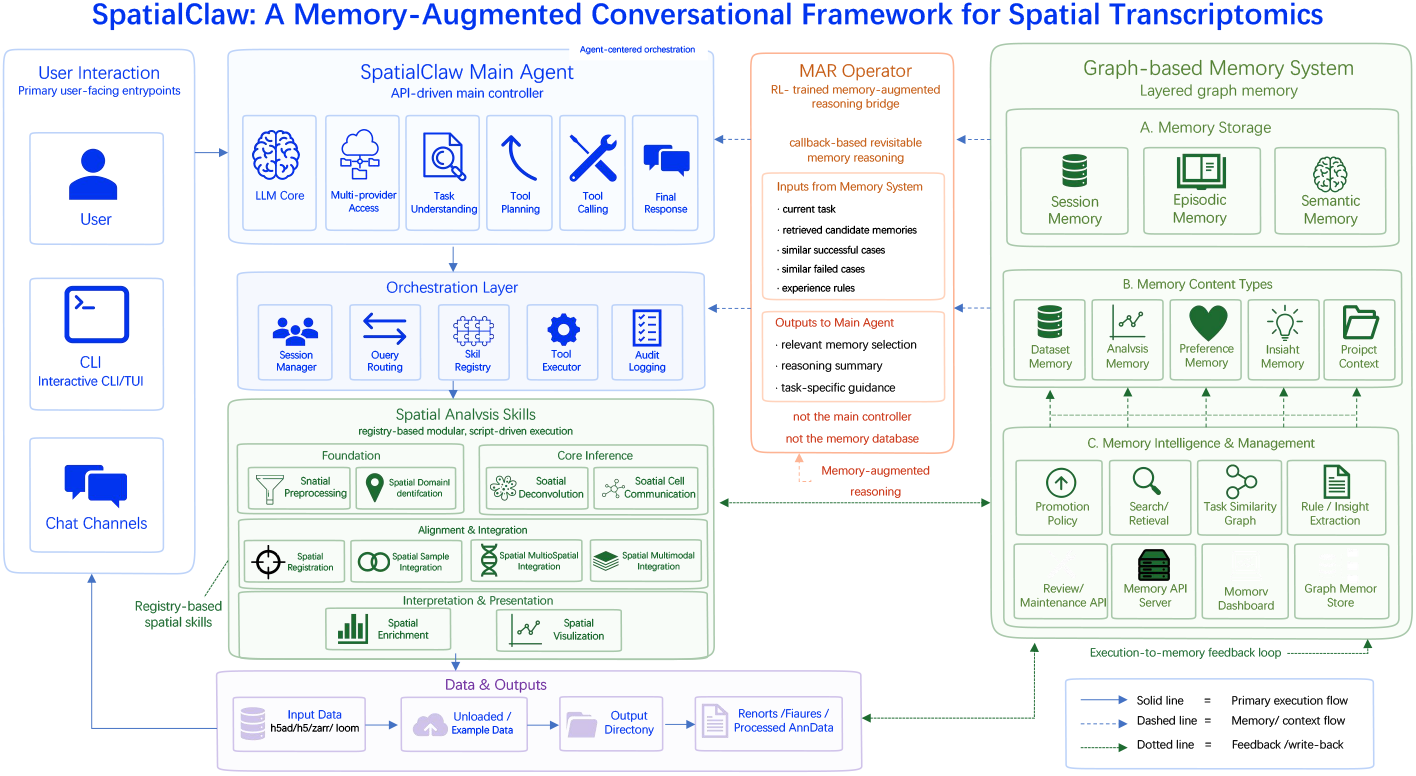
Overview of the SpatialClaw autonomous ecosystem. SpatialClaw integrates a conversational LLM agent, a modular spatial-analysis skill library, and a graph-based persistent memory system. User queries arrive through three entry points (CLI/TUI, chat channels, and API) and are processed by the SpatialClaw Main Agent, which handles task understanding, tool planning, and response generation via the Orchestration Layer (session management, query routing, skill registry, tool execution, and audit logging). The Spatial Analysis Skills catalog organizes 30 registered skills into four functional tiers—Foundation, Core Inference, Alignment & Integration, and Interpretation & Presentation—and accepts h5ad/h5/zarr/loom input, writing structured outputs to a timestamped directory. A Memory-Augmented Reasoning (MAR) Operator bridges the agent and the Graph-based Memory System: for each task, it retrieves candidate memories (similar successful and failed cases, distilled experience rules) and returns a reasoning summary and task-specific guidance to the main agent. The memory system stores three hierarchical layers—Session, Episodic, and Semantic Memory—populated by five typed objects (Dataset, Analysis, Preference, Insight, and Project Context). Memory intelligence is managed through a Promotion Policy, Search/Retrieval, a Task Similarity Graph, and a Rule/Insight Extraction loop, with all AI-induced mutations exposed through a Review/Maintenance API and a Memory Dashboard. Solid arrows: primary execution flow; dashed arrows: memory/context flow; dotted arrows: feedback write-back.

#### Agent and orchestration

The SpatialClaw Main Agent serves as the sole control point for all user-facing interactions, handling task understanding, tool planning, and response generation. Supporting it, the Orchestration Layer manages session identity, skill selection, catalog discovery, subprocess execution, and immutable invocation records. Crucially, all three user-facing entry points—interactive CLI/TUI, chat adapters (Telegram and Feishu), and direct API—converge on the same Orchestration Layer, so execution semantics are identical regardless of interface.

#### Skill library

The Spatial Analysis Skills catalog currently ships 30 registered skills (29 spatialomics, 1 orchestrator), organized into four functional tiers: Foundation (preprocessing, spatial domain identification), Core Inference (deconvolution, Cell–cell communication), Alignment & Integration (multi-sample integration, cross-modal fusion, registration), and Interpretation & Presentation (enrichment, visualization). Each skill is a self-contained directory exposing a uniform command-line interface and is invoked as an isolated subprocess, preventing heavyweight dependencies from bloating the interactive process. Every skill writes a fixed output contract—a Markdown narrative report, a machine-readable result envelope, an updated AnnData when applicable, and a reproducibility bundle capturing the exact command, environment, and input checksums.

#### Memory system and MAR Operator

The Graph-based Memory System stores five typed memory objects across three hierarchical layers (Session, Episodic, and Semantic), governed by a deterministic promotion policy and augmented by a Memory-Augmented Reasoning (MAR) Operator that synthesizes retrieved experiences into task-specific guidance; the architecture and empirical evaluation are described in §2.3.

### 2.2 Comprehensive tool coverage across spatial-omics tasks

We evaluated SpatialClaw’s capability profile against 12 representative platforms using a structured relative-strength matrix covering 15 functional dimensions (Fig. 2). The evaluation framework spans three domains: Core Tasks (Prep, Structure, Annot/Comp, Stats/SVG, Compar/Enrich), Extended Modalities (AdvInf, Integr/Reg, Pred/Imp, Histol/WSI, X-omics), and Platform capabilities (Memory, Trace, Viewer, Closure, Local). Only explicitly supported and documented capabilities are marked as directional; all others are conservatively marked as “≈”.

**Figure 2:**
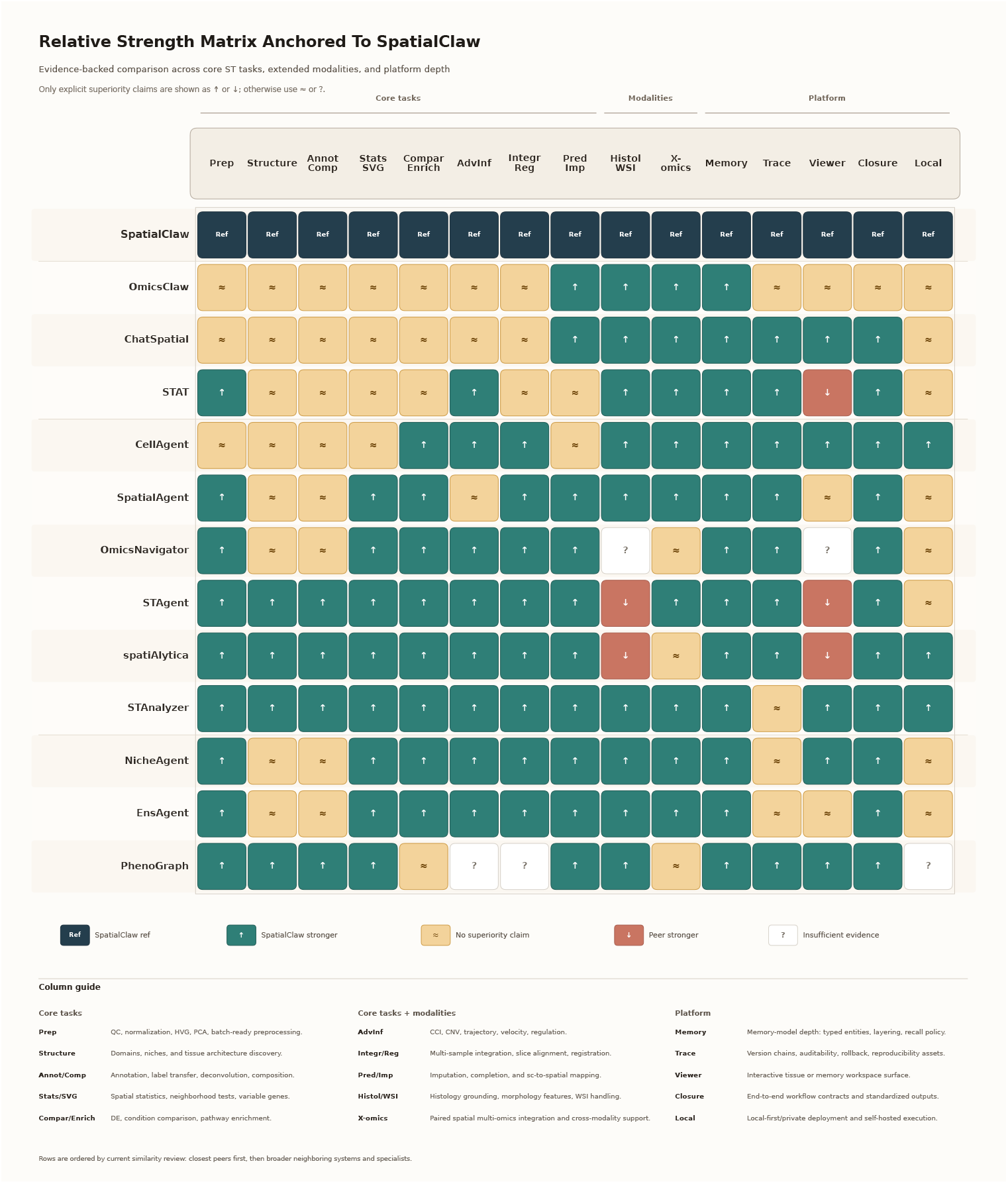
Relative capability matrix of SpatialClaw against representative spatial-omics agent platforms. Rows represent platforms, columns represent functional capabilities, and Spatial-Claw serves as the reference baseline (dark “Ref” cells). Columns span Core Tasks (Prep, Structure, Annot/Comp, Stats/SVG, Compar/Enrich), Extended Modalities (AdvInf, Integr/Reg, Pred/Imp, Histol/WSI, X-omics), and Platform capabilities (Memory, Trace, Viewer, Closure, Local); full definitions are provided in the column guide at figure bottom. Cell color and label indicate the direction of any demonstrated capability difference: dark teal “↑” cells indicate SpatialClaw is stronger; orange “↑” cells indicate the peer is stronger; sand “≈” cells indicate no superiority claim; white “?” cells indicate insufficient evidence. Only claims supported by documented evidence are shown as directional; all other comparisons are conservatively marked “≈”. Platforms are ordered by similarity to SpatialClaw, with closest peers (OmicClaw, ChatSpatial) listed first.

Across Core Tasks, SpatialClaw’s 29 dedicated spatial-omics skills provide reference-level coverage from preprocessing through enrichment without requiring the user to leave the platform. In Extended Modalities, integrations with Tangram, Stereoscope, SpaGCN, STAGATE, GraphST, BANKSY, SpatialGlue, Harmony, Scanorama, DESeq2, CellPhoneDB, and CellChat collectively cover deconvolution, multi-sample integration, histology, and cross-modal fusion—capabilities that most platforms address only partially. In the Platform domain, the graph-based memory subsystem, changeset-backed audit trail, and reproducibility bundle underpin the Memory, Trace, and Closure columns.

OmicClaw and ChatSpatial, the two most architecturally similar platforms, achieve parity on Core Tasks but trail SpatialClaw on Memory, Trace, and Closure, lacking the typed, layered, graph-backed memory and changeset audit trail that SpatialClaw provides. SpatialAgent’s memory–plan–execute loop represents a step beyond purely stateless designs but does not implement versioned graph storage or deterministic promotion-based episodic-to-semantic elevation. Specialist platforms (STAT, STAgent, spatiAlytica, STAnalyzer) achieve strong performance on Core Tasks consistent with their published focus, yet uniformly lack documented equivalents for Memory, Trace, and Closure. Across the entire matrix, these two columns emerge as the most consistent differentiators: no peer platform implements typed, layered, graph-backed memory with semantic retrieval and rollback comparable to SpatialClaw’s, and standardized end-to-end output contracts remain absent from most systems. Together, these gaps explain why existing agents—despite strong analytical capabilities—require users to re-upload data and reconfigure parameters at the start of every new session.

### 2.3 Persistent memory enables continuity across sessions

A central design goal of SpatialClaw is to maintain analytical continuity across sessions and across extended multi-step workflows without requiring users to re-upload data or re-specify parameters at each interaction. To operationalize this, SpatialClaw implements a graph-based persistent memory subsystem and a Memory-Augmented Reasoning (MAR) Operator, whose architecture and empirical benefit are described in this section.

#### Memory architecture (Fig. 3a)

The memory subsystem operates across three levels. At the storage level, raw records generated during agent execution—user queries, tool calls, results, and error events—are parsed into structured memory triplets of the form (Entity, Relation, Entity) and written to a persistent graph database (Neo4j) that supports ACID transactions, versioning, and time-stamping across sessions. At the representational level, the memory graph maintains two parallel layers: an *Episodic Memory* layer that stores short-term, session-specific records (recent queries, tool calls, messages, errors, and results as a sequential event chain), and a *Semantic Memory* layer that stores long-term reusable knowledge organized around stable spatial-omics entities (Project, Sample, Cell Type, Region, Slide). Bridging these two layers are five typed memory objects—DatasetMemory, AnalysisMemory, ProjectContextMemory, InsightMemory, and PreferenceMemory—each capturing a distinct category of analytical state. A *Memory Promotion Policy* governs the elevation of episodic records to semantic memory based on four criteria: stability, confidence, relevance, and evidence; only records that meet these criteria are durably retained, keeping the semantic layer precise rather than exhaustive. At the operational level, five graph operations maintain the memory store over time: Write (entity extraction, node/edge creation, metadata attachment), Read (subgraph retrieval, multi-hop reasoning, hybrid vector+graph search), Update (node/edge upsert, version control, conflict resolution), Summarize (community summary, key fact extraction, abstraction and pruning), and GC (TTL-based ephemeral node removal, orphan cleanup, archival). The MAR Operator sits between the memory store and the main agent, implementing a five-step per-query loop: (1) Understand & Retrieve— retrieve the relevant subgraph via multi-hop traversal; (2) Plan & Reason—leverage retrieved memory to decompose tasks and select tools; (3) Execute—invoke the selected skill; (4) Write Back—update the memory graph with new records and relations; and (5) Summarize & Persist—compress and persist the session for future retrieval. This loop is designed to be iterative across sessions, enabling the agent to accumulate and reuse analytical experience over long-running projects.

#### Memory benchmark design

To quantify the benefit of the persistent memory subsystem and the MAR Operator, we designed a controlled benchmark spanning three memory-sensitive scenarios, each evaluated across the same 10 registered spatial-omics skills (spatial-preprocessing, spatial-domain-identification, spatial-deconvolution, spatial-cell-communication, spatial-registration, spatial-multi-sample-integration, spatial-omics-integrate, spatial-modality-integrate, spatial-enrichment, and spatial-visualization). Three system configurations were compared: *baseline memory* (a standard LLM without SpatialClaw’s structured memory), *spatialclaw memory* (SpatialClaw’s graph-backed typed memory without MAR, `SPATIALCLAW_REMEM_ENABLED=false`), and *spatialclaw memory + MAR* (`SPATIALCLAW_REMEM_ENABLED=true`, `SPATIALCLAW_REMEM_INVOKE_MODE=always`). Each skill was evaluated in an independent temporary database to prevent cross-skill memory contamination. Scores were computed per skill according to skill-specific rubrics covering parameter recall, detail retention, and continuation fidelity; per-skill scores were summed to yield a total for each scenario. Tool executors were stubbed throughout, so the benchmark measures memory quality rather than analysis correctness.

The three scenarios operationalize distinct memory demands. *Resumed-session* (Fig. 3b; maximum score 118) tests cross-session continuity: a prior session is saved, the process state is cleared, the session is reloaded, and the agent is asked to recall and continue the prior workflow through recall, continuation, and control prompts. *Multi-turn* (Fig. 3c; maximum score 138) tests same-session continuity: within a single live conversation, four sequential prompts (recall, detail, continuation, control) probe whether the agent sustains coherent memory of the workflow as the dialogue progresses. *Tool-continuation* (Fig. 3d; maximum score 168) tests parameter-level continuity: after a prior analysis, the agent is asked to resume the workflow and invoke the next tool directly, and scoring checks whether historically successful parameters are correctly propagated into the real `spatialclaw(…)` tool call.

**Figure 3:**
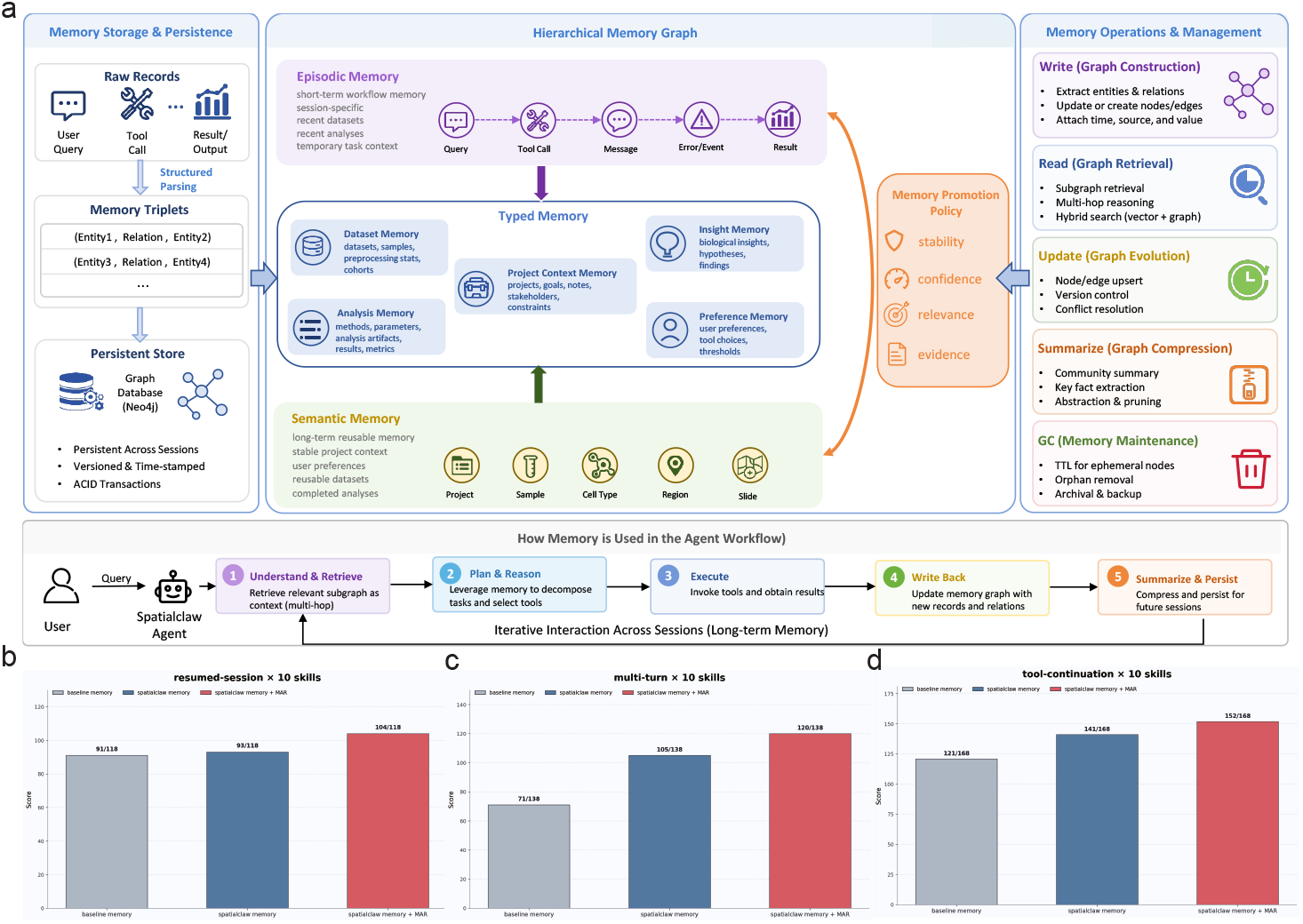
The SpatialClaw persistent memory subsystem and its benchmark evaluation. **(a)** Architecture of the memory subsystem and the Memory-Augmented Reasoning (MAR) Operator. Left: raw records (user queries, tool calls, results) are parsed into structured memory triplets and written to a persistent graph database (Neo4j) with ACID transactions, versioning, and time-stamping. Center: the Hierarchical Memory Graph maintains an Episodic Memory layer (short-term, session-specific event chains) and a Semantic Memory layer (long-term reusable knowledge organized around spatial-omics entities: Project, Sample, Cell Type, Region, Slide), connected by five typed memory objects (Dataset, Analysis, Project Context, Insight, and Preference Memory). A Memory Promotion Policy evaluates episodic records on four criteria (stability, confidence, relevance, evidence) before elevating them to the semantic layer. Right: five graph operations (Write, Read, Update, Summarize, GC) maintain the memory store over time. Bottom: the five-step per-query agent workflow—Understand & Retrieve → Plan & Reason → Execute → Write Back → Summarize & Persist—is designed for iterative use across sessions, enabling long-term memory accumulation. **(b–d)** Benchmark evaluation of three memory configurations—baseline memory, SpatialClaw memory, and SpatialClaw memory + MAR—across three memory-sensitive scenarios, each evaluated over 10 spatial-omics skills. **(b)** Resumed-session scenario (maximum score 118): a prior session is saved, process state is cleared, and the session is reloaded; scores measure cross-session recall and continuation fidelity. Scores: baseline 91, SpatialClaw memory 93, +MAR 104. **(c)** Multi-turn scenario (maximum score 138): within a single live session, four sequential prompts (recall, detail, continuation, control) test sustained memory coherence across conversational turns. Scores: baseline 71, SpatialClaw memory 105, +MAR 120. **(d)** Tool-continuation scenario (maximum score 168): the agent is asked to resume a prior workflow and directly invoke the next tool; scoring checks whether historically successful parameters are correctly propagated into the tool call. Scores: baseline 121, SpatialClaw memory 141, +MAR 152. Tool executors were stubbed throughout; benchmark measures memory quality rather than analysis correctness.

#### Persistent memory substantially outperforms baseline across all three scenarios

Replacing baseline memory with SpatialClaw’s structured graph memory alone yields consistent gains: in the resumed-session scenario the score rises from 91 to 93 (out of 118); in the multi-turn scenario from 71 to 105 (out of 138); and in the tool-continuation scenario from 121 to 141 (out of 168). The multi-turn scenario shows the largest absolute gain (+34 points), reflecting that a flat context buffer degrades rapidly under sequential multi-turn prompting whereas a structured graph store maintains indexed access to prior workflow state regardless of conversation length.

#### The MAR Operator provides consistent additional benefit on top of structured memory

Adding MAR to SpatialClaw’s memory further improves scores to 104/118 in resumed-session (+11 over memory alone), 120/138 in multi-turn (+15), and 152/168 in tool-continuation (+11). The multi-turn scenario again shows the largest MAR gain, consistent with MAR’s role as a reasoning bridge that actively synthesizes retrieved memories into task-specific guidance across successive conversational turns rather than passively surfacing stored records. Skill-level analysis reveals that MAR benefits are concentrated in continuation-heavy skills: in tool-continuation, spatial-domain-identification improves from 13/15 to 15/15, spatial-deconvolution from 17/21 to 21/21, and spatial-registration from 15/17 to 17/17, with the gains attributable to more reliable propagation of structured parameters such as `resolution`, `reference_path`, `cell_type_key`, and `reference_slice` into downstream tool calls. In contrast, skills where baseline memory already provides sufficient context (e.g., spatial-preprocessing, spatial-multi-sample-integration) show little or no additional gain from MAR, indicating that the operator adds value where retrieval quality alone is insufficient to resolve parameter ambiguity. One skill (spatial-preprocessing in resumed-session) showed a minor score decrease under MAR (7*/*8 versus 8*/*8), traced to a sentinel memory leakage in the control prompt; this edge case highlights that over-retrieval of irrelevant prior context remains a limitation of the current promotion policy.

Taken together, these results demonstrate that SpatialClaw’s graph-based persistent memory and MAR Operator jointly address the cross-session and multi-turn continuity gaps that constrain existing spatial-omics agents, enabling users to resume prior workflows, continue multi-step analyses, and propagate historical parameters into new tool calls without manual reconfiguration.

### 2.4 End-to-end analysis of a high-heterogeneity breast cancer spatial omics cohort

To evaluate SpatialClaw’s ability to orchestrate a complete spatial-omics workflow on real, complex data, we applied it to a publicly available human triple-negative breast cancer (TNBC) cohort [25] (GEO: GSE213688), profiled by 10x Genomics Visium. The cohort spans samples M1–M16, with M12 absent, yielding 15 usable sections that collectively capture diverse tissue microenvironments including tumor, stroma, adipose, and inflammatory regions. Our aim was not to propose new mechanistic findings on this dataset, but to demonstrate that SpatialClaw can stably orchestrate—through natural-language instructions in three conversational turns—the full analytical chain from multi-section preprocessing through spatial domain identification, marker extraction, functional enrichment, semantic annotation, and inter-domain communication analysis, while producing auditable, standardized outputs at every step.

#### Cohort ingestion and preprocessing quality control (Fig. 4A–B)

The 15 sections were ingested as raw `h5ad` files through a single session-initialization command (Fig. 4A). SpatialClaw’s preprocessing skill executed spot-level QC filtering, library-size normalization, log transformation, highly variable gene selection, PCA, and neighborhood graph construction in a single automated pass, propagating the processed `AnnData` object to the session’s primary data path for all downstream skills without requiring redundant uploads. Violin plots of per-spot detected gene count (*n_genes_by_counts*) and total UMI count (*total_counts*) across the pooled cohort confirmed that samples passed QC thresholds and were suitable for downstream spatial analysis (Fig. 4B).

**Figure 4:**
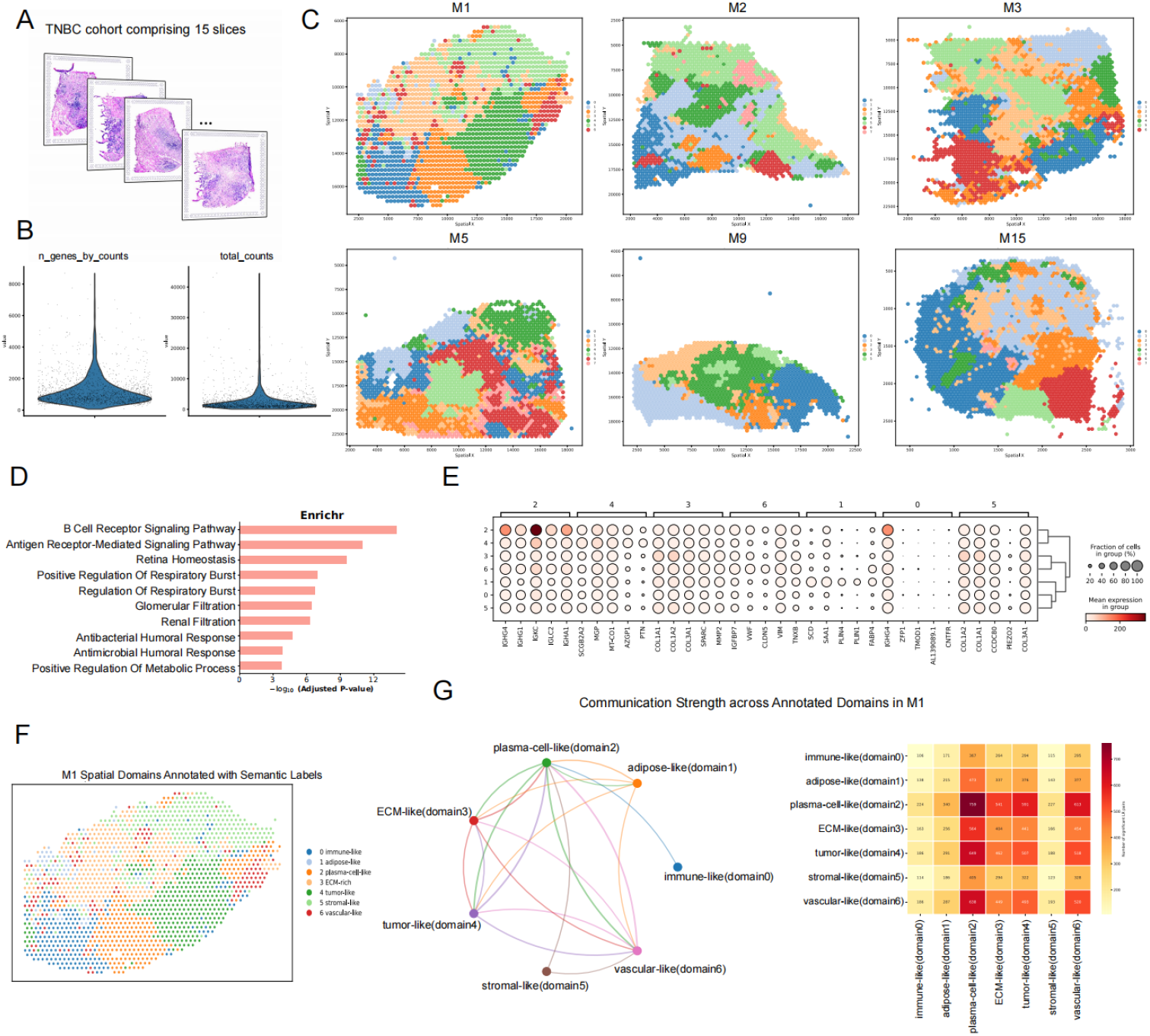
End-to-end spatial omics analysis of a human TNBC cohort through Spatial-Claw (GSE213688). **(A)** Schematic of the TNBC cohort comprising 15 10x Visium tissue sections (M1–M16, with M12 absent), illustrating morphological diversity across samples. **(B)** Preprocessing quality control metrics aggregated across the cohort, showing per-spot distributions of detected gene count (*n_genes_by_counts*) and total UMI count (*total_counts*), confirming data quality prior to downstream analysis. **(C)** Spatial domain identification results for six representative sections (M1, M2, M3, M5, M9, M15) using BANKSY. Capture spots are colored by domain label and overlaid on physical tissue coordinates. Between 4 and 9 domains were identified per section under a unified parameter setting, reflecting genuine inter-section heterogeneity in tissue architecture. **(D)** Domain-resolved GO enrichment results for a representative spatial domain in section M1 (Enrichr, ranked by − log_10_(adjusted *p*-value)). The top enriched terms include B Cell Receptor Signaling Pathway (GO:0050853), Antigen Receptor-Mediated Signaling Pathway (GO:0050851), and immune-effector processes (respiratory burst, humoral immune response), supporting functional annotation of this domain as an immune-active region. **(E)** Per-domain marker gene dot plot for M1. Dot size encodes the fraction of spots expressing each gene within the domain; color encodes mean expression level. Domain-specific molecular signatures support downstream semantic annotation. **(F)** Semantic annotation of M1’s seven spatial domains based on marker genes and domain-resolved enrichment results. Domains are labeled as *immune-like* (0), *adipose-like* (1), *plasma-cell-like* (2), *ECM-like* (3), *tumor-like* (4), *stromal-like* (5), and *vascular-like* (6), and mapped back to spatial coordinates. **(G)** Inter-domain cell–cell communication analysis for M1 using semantic domain labels as input (LIANA; 16,753 ligand–receptor pairs tested, 3,087 significant). Left: chord diagram in which edge thickness encodes aggregated communication strength between sender–receiver domain pairs. Right: communication-strength heatmap across all 7 *×* 7 domain pairs. High-scoring interactions are concentrated between *ECM-like*/*stromal-like* domains and *adipose-like, tumor-like*, and *vascular-like* domains, with representative pairs *COL1A1* /*COL1A2* → *CD36, COL1A1* /*COL1A2* → *CD44, SPARC* → *ENG*, and *COL1A1* → *ITGA5*, consistent with extracellular-matrix-associated inter-domain crosstalk.

#### Spatial domain identification across representative sections (Fig. 4C)

Spatial domain identification was performed on all 15 sections using BANKSY, which extends graph-based clustering by incorporating a neighborhood expression embedding that captures local microenvironmental context beyond individual spot identity. Fig. 4C shows results for six representative sections (M1, M2, M3, M5, M9, and M15). The identified domains ranged from 4 to 9 per section under a unified parameter setting, reflecting genuine differences in tissue architecture and transcriptional heterogeneity rather than parameter instability. Sections with morphologically compact tumor regions (M1, M2) exhibited well-delineated domains, while sections with more dispersed tissue organization (M5, M9, M15) showed finer spatial mosaics consistent with their histological complexity. Across all 15 sections, SpatialClaw successfully completed preprocessing and domain identification, generating for each sample a structured output bundle comprising a processed `h5ad` file, spatial domain visualizations, `report.md`, `result.json`, and a reproducibility record capturing the exact command and environment.

#### Domain marker extraction and functional enrichment (Fig. 4D–E)

To move beyond anonymous domain labels, we directed SpatialClaw to extract per-domain marker genes for section M1 and to perform domain-resolved GO enrichment analysis using Enrichr, constructing a functional bridge between unsupervised spatial partitioning and interpretable tissue states. The domain-resolved enrichment results (Fig. 4D) show that the top-ranked domain is strongly enriched for immune-effector processes: the two leading terms are B Cell Receptor Signaling Pathway (GO:0050853) and Antigen Receptor-Mediated Signaling Pathway (GO:0050851), followed by respiratory burst regulation (GO:0060267, GO:0060263), humoral immune response terms (GO:0019731, GO:0019730), and metabolic process regulation (GO:0009893). The per-domain marker gene dot plot (Fig. 4E) further characterizes domain-specific molecular signatures: domains 2 and 4 are distinguished by immunoglobulin-related genes (*IGKC, IGLC2, IGLC1*) and collagen/matrix genes (*COL1A1, COL1A2, COL3A1, SPARC*), respectively; domain 1 expresses adipocyte markers (*FABP4, PLIN1*); domain 6 shows vascular markers (*VWF, CLDN5*); and domain 5 is characterized by stromal genes alongside *ZFP1* and *AL139089*.*1*. Together, these marker and enrichment profiles provide the molecular evidence base for semantic annotation in the next step.

#### Semantic domain annotation (Fig. 4F)

Based on the domain-resolved marker genes and GO enrichment results, SpatialClaw assigned functional semantic labels to M1’s seven spatial domains: domain 0 as *immune-like*, domain 1 as *adipose-like*, domain 2 as *plasma-cell-like*, domain 3 as *ECM-like*, domain 4 as *tumor-like*, domain 5 as *stromal-like*, and domain 6 as *vascular-like* (Fig. 4F). The *plasma-cell-like* and *immune-like* domains are jointly supported by enrichment in B cell receptor signaling and humoral immune response, consistent with an antibody-secreting immune infiltrate. The *adipose-like* domain is supported by lipid metabolism and fatty acid transport terms together with canonical adipocyte markers (*PLIN1, FABP4, LPL*). The *ECM-like* and *stromal-like* domains both show enrichment in extracellular matrix organization and collagen fibril organization, with the former reflecting denser matrix remodeling and the latter a more fibroblast-like interstitial state. The *vascular-like* domain is supported by vasculogenesis and angiogenesis-related pathways alongside *VWF* and *PECAM1* expression, and the *tumor-like* domain exhibits a keratin-associated epithelial program characteristic of carcinoma tissue. This step demonstrates SpatialClaw’s capacity to advance from unsupervised spatial partitioning to histologically and functionally interpretable spatial states through a single additional conversational instruction.

#### Inter-domain cell–cell communication (Fig. 4G)

Using the semantic domain labels as input, we applied SpatialClaw’s LIANA-based cell–cell communication skill to M1. Of 16,753 ligand– receptor pairs tested, 3,087 reached statistical significance. The chord diagram and communication-strength heatmap (Fig. 4G) reveal that the highest-scoring interactions are concentrated between the *ECM-like* and *stromal-like* domains and the *adipose-like, tumor-like*, and *vascular-like* domains. Representative high-scoring pairs include *COL1A1* /*COL1A2* → *CD36, COL1A1* /*COL1A2* → *CD44, COL1A1* /*COL1A2* → *CD93, SPARC* → *ENG*, and *COL1A1* → *ITGA5*. This pattern is consistent with a strong extracellular-matrix-associated inter-domain crosstalk signature characteristic of this high-stroma TNBC cohort. The communication results should be interpreted as spatially resolved ligand–receptor co-expression evidence rather than direct assertions of intercellular mechanisms; their primary value here is to demonstrate that SpatialClaw seamlessly propagates upstream semantic domain labels into the downstream communication module, producing histologically interpretable output without any manual data reformatting between steps.

Taken together, this case study demonstrates that SpatialClaw can stably orchestrate—in three conversational turns and with zero direct scripting—a multi-step spatial-omics workflow spanning preprocessing, spatial domain identification, marker extraction, functional enrichment, semantic annotation, and inter-domain communication analysis across a 15-section cohort. The standardized output bundle generated at each step—comprising figures, tables, structured result objects, and reproducibility records—illustrates that SpatialClaw’s advantage lies not only in executing individual analysis tasks but in organizing multiple inter-dependent steps into a traceable, reproducible, and interpretable end-to-end pipeline.

## 3 Discussion

We have presented SpatialClaw, an end-to-end LLM-agent autonomous ecosystem that unifies spatialomics analysis under a single conversational interface and a persistent, structured memory subsystem. The autonomous ecosystem architecture (Fig. 1) reduces every user interaction—across CLI, TUI, and chat frontends—to a single execution path in which a natural-language query is routed to one of 30 registered skills, its arguments are normalized against a declarative schema, and its outputs are written through a fixed report contract. The skill catalog (Fig. 2) covers the major analytical axes of spatial omics, from preprocessing and spatial-domain identification to deconvolution, spatially variable gene detection, cell–cell communication, multi-sample integration, and cross-modality fusion, and is reachable through a single conversational layer rather than through manual tool stitching. The memory subsystem (Fig. 3) further substantiates that, with persistent memory enabled, multi-turn trajectories require fewer redundant uploads and fewer user turns to reach a valid first result, and analyses interrupted in one session can be resumed in another with full lineage. End-to-end natural-language reproduction of a published study (Fig. 4) demonstrates that the combination of broad tool coverage and persistent memory translates into a complete, traceable analytical pipeline delivered in three conversational turns.

Beyond these aggregate numbers, we believe the more durable contribution of SpatialClaw lies in two design positions that we hope will outlast the specific implementation. The first is that spatial omics is now methodologically diverse enough to warrant a domain-dedicated agent rather than a general-purpose bioinformatics assistant: the heterogeneity of platforms, the co-existence of single-slide, multi-slide, and multi-modal workflows, and the rapidly expanding set of method-specific parameter spaces are difficult to harmonize without a curated registry, declarative argument schemas, and a routing layer that is aware of the field’s vocabulary. SpatialClaw demonstrates that this harmonization can be achieved without sacrificing extensibility: new skills are added as self-contained directories with a uniform CLI contract, and the registry’s lazy-loading discovery keeps cold-start latency bounded as the catalog grows. The second position is that agent memory should be a first-class, structured, and auditable substrate rather than an ephemeral context buffer or a flat vector store [18, 19]. The four-entity graph model (Node/Memory/Edge/Path) cleanly separates entity identity, content versioning, structural relationship, and external addressing; the parallel episodic and semantic layers, governed by a deterministic promotion policy, distinguish raw session traces from durable transferable knowledge [21]; and the changeset-based audit trail makes every AI-induced mutation inspectable and reversible. Together these decisions enable upload-once analyse-many workflows, project-level continuity over days and across collaborators, and cross-project reuse of confirmed insights, while ensuring that the raw expression matrices themselves never leave the user’s environment.

Several limitations of the current system warrant explicit mention. At the architectural level, SpatialClaw orchestrates a single agent over a serial execution path; complex workflows that would benefit from parallel exploration, separation of planning and execution, or independent auditing roles must currently be sequenced manually by the user. At the memory level, the promotion policy, the Jaccard threshold for similarity edges, the insight score initialization, and the rule-set cap are all empirical constants tuned for spatial omics; their portability to other domains is plausible but not yet validated. The insight-extraction loop, in which the LLM is asked to Add, Edit, Remove, or Agree on accumulated rules, is itself susceptible to model hallucination [20]; we mitigate this through score-based decay, periodic deduplication, and per-cluster rule-set caps, but the underlying dependence on LLM judgement remains. Privacy is enforced structurally—absolute paths and multi-line payloads are rejected by validators, and only metadata is persisted—but semantic-level filtering of free-text content (for example, inadvertent inclusion of patient identifiers in a user comment) is not yet performed. The user study, while controlled and blinded, was conducted on a finite task suite with a finite participant pool, and its baselines are limited by what existing platforms can natively execute on spatial-omics workloads.

These limitations chart a natural research agenda. The most immediate direction is to evolve the single-agent execution loop into a multi-agent topology in which planning, execution, and auditing are separated, enabling parallel exploration of parameter spaces and proactive proposal of follow-up analyses or controlled experiments rather than purely reactive response to user queries. A second direction is to deepen multi-modal integration: with the memory graph as a cross-modal hub, hematoxylin-and-eosin and immunofluorescence images, whole-slide pathology, single-cell and bulk transcriptomes can be aligned as first-class nodes whose insights become reachable across modalities. A third direction is to open the skill registry as a community-curated catalog, so that new spatial-omics methods can be contributed and versioned without forking the core platform, and to develop visualization, inspection, and controlled-editing tools for the memory graph itself so that users can examine, audit, and correct the rules the agent has learned—an interpretability layer for which the existing changeset and rollback machinery already provides a foundation. Taken together, we view SpatialClaw not as a finished tool but as a substrate on which methods, memory, and users can co-evolve toward traceable, reproducible, and increasingly autonomous spatial-omics research.

## 4 Methods

The SpatialClaw implementation is a Python package that exposes a single LLM agent over a curated skill catalog and a graph-backed persistent memory system. We describe the platform in three parts: §4.1 specifies the end-to-end execution pipeline—how a user query is parsed, routed, normalized, and run as a subprocess; §4.2 specifies the memory mechanism—its data model, storage path, retrieval, and auditability; §4.3 specifies what is required to reproduce a run and how reproducibility is enforced inside the platform itself.

### 4.1 End-to-end pipeline

SpatialClaw is structured as a thin domain-agnostic framework (`spatialclaw/`) on top of a curated skill catalog (`skills/`). The framework provides exactly one execution path—natural-language query → router → argument normalization → subprocess → report and memory write—and that path is shared by every entry point, so behaviour at the CLI, in the TUI, and through chat adapters is identical.

#### 4.1.1 Skill abstraction and the registry

Every analysis task in SpatialClaw is implemented as a *skill* : a self-contained directory under `skills/<domain>/<skill-name>/` containing (i) a YAML-fronted Markdown specification `SKILL.md` declaring identifier, version, dependencies, and trigger keywords; (ii) one Python entry-point script that exposes a uniform CLI (`–input`, `–output`, plus skill-specific flags); and (iii) an optional test suite. The current release ships **30 skills** (29 spatial-omics skills covering preprocessing, domain identification, deconvolution, statistics, differential expression, spatially variable genes, cell–cell communication, CNV inference, multi-sample integration, cross-modality integration, registration, trajectory inference, velocity, enrichment, histology, morphology, WSI processing, oncology, TLS detection, target discovery, niche analysis, regulation, cell annotation, sc2spatial mapping, translation, and visualization; plus 1 orchestrator skill providing query routing). The complete enumeration is in Supplementary Table S1.

Skill discovery is the responsibility of an `AnalysisRegistry` singleton in `spatialclaw.core. registry`. The registry combines two mechanisms: a curated table of production-grade skills with hard-coded argument schemas, trigger keywords, and parameter validation rules (guaranteeing stable behaviour for skills that have been audited), and a dynamic filesystem scan that picks up any additional skill directory conforming to the layout above. A `LazySkillMetadata` proxy parses only the YAML frontmatter of `SKILL.md` on demand and defers full module import until first invocation, keeping interactive cold start under one second even with the full catalog registered.

#### 4.1.2 Argument normalization layer

LLM-emitted tool calls routinely exhibit lexical variation in parameter names and values—`save_dir` versus `output_dir`, `slide-seqv2` versus `Slide-seqV2`, `spatial-glue` versus `spatialglue`. Left untreated, this single failure mode is responsible for the majority of broken tool calls in LLM-driven pipelines. SpatialClaw resolves it before any subprocess is spawned, through a two-stage normalization layer implemented by `normalize_skill_llm_args` and `build_skill_llm_cli_args` in the registry. A *global* schema harmonizes universal arguments (`output_dir`, `method`, `n_epochs`, `device`, `platform`) across all skills, and per-skill *overrides* declared in the skill’s `llm_argument_schema` block specify allowed values, value aliases, numeric bounds, and a `drop_invalid` flag that controls whether out-of-spec values are silently dropped or passed through. Normalization performs case-insensitive alias resolution, clamping against numeric bounds, and rejection of values that fall outside the `allowed_ values` whitelist when `drop_invalid` is set. Only canonical arguments whose CLI flag is in the skill’s `allowed_extra_flags` whitelist are forwarded to the subprocess, ensuring that the LLM cannot inject arbitrary flags into a skill’s command line.

#### 4.1.3 Routing

Mapping a user query to one of the 30 registered skills is delegated to a unified router (`route_query_ unified` in `spatialclaw.routing.router`) supporting three modes. *Keyword mode* matches the query against the `trigger_keywords` declared in each `SKILL.md`, ranking candidate skills by total matched-keyword length and returning a confidence score min(1.0, matched_length*/*20); it is fully deterministic and requires no API call. *LLM mode* assembles skill names and one-line descriptions into a prompt and asks the configured LLM to return a JSON object `{“skill”: …, “confidence”: …}` with the confidence clamped to [0, 1] and the skill validated against the registered set; invalid skills cause a fallback to the null result. The routing module is provider-agnostic: a `_resolve_llm_config` helper reads `LLM_PROVIDER` or auto-detects from environment variables, and the implementation supports twelve LLM backends (Anthropic Claude, OpenAI GPT, Google Gemini, DeepSeek, Qwen via DashScope, GLM via Zhipu, Doubao via Volcengine, NVIDIA NIM, SiliconFlow, OpenRouter, Ollama for local hosting, and a custom OpenAI-compatible endpoint), each with a fixed preset of base URL, default model, and API-key environment variable. *Hybrid mode* (the default for interactive sessions) issues the keyword router first and falls back to the LLM router only when keyword confidence falls below a configurable threshold (default 0.5). The selected skill, together with the LLM-emitted arguments coerced through the normalization layer of §4.1.2, is then passed to the execution layer.

#### 4.1.4 Execution and report contract

Skill execution is performed by `run_skill` in `spatialclaw.py`. The function resolves the input path (absolute path to a data file, a previously persisted `AnalysisSession`, or the skill’s built-in `–demo` fixture), allocates a timestamped output directory under `output/<skill>_<timestamp>/`, and constructs the command line `[python, <script>, –input <path>, –output <dir>, <extra flags>]`. Required-input rules (`required_extra_inputs`) and alternative-input modes (`allows_alternative_inputs`) declared in the registry are checked before launch. The skill is then invoked through `subprocess.run` with `capture_output=True` and `PYTHONPATH` pre-pended, so the parent process never imports the skill’s heavyweight scientific dependencies (Scanpy, Squidpy, PyTorch, etc.) and remains responsive in interactive sessions.

Every skill writes to its output directory a fixed contract (`spatialclaw.common.report`): a Mark-down narrative report `report.md`, a machine-readable envelope `result.json` containing skill identifier, version, completion timestamp, input SHA-256 checksum, summary, and structured data, and—when the skill is marked `saves_h5ad` in the registry—an updated `processed.h5ad` that downstream skills can chain from. Tabular results live under `tables/`, plots under `figures/`, and a `reproducibility/` sub-directory captures the exact command, environment specification, and SHA-256 checksums of all inputs. Both `report.md` and the assistant’s chat output append the standard SpatialClaw disclaimer that the platform is a research tool and not a clinical device.

To support upload-once, analyze-many workflows, the `AnalysisSession` object in `spatialclaw. common.session` persists session metadata (input file, dataset checksum, species, data type, domain) and accumulated skill results to a JSON file. Subsequent skill calls within the same session resolve their input from the session’s `primary_data_path`, eliminating redundant uploads across multi-step analyses.

#### 4.1.5 Frontends and dependency management

Three user-facing frontends share this single backend: a prompt-toolkit REPL with slash commands and Rich-rendered output (`spatialclaw interactive`), a full-screen Textual TUI (`spatialclaw tui`), and chat adapters for Telegram and Feishu (`bot/`). All three resolve to the same `run_skill` entry point, so the execution semantics described above are independent of the surface. A research-mode pipeline in `spatialclaw.agents` optionally orchestrates idea-to-report workflows over multiple skills using LangGraph and DeepAgents, and a pluggable ReMemR1 reasoning operator (`spatialclaw.agents.remem_operator`) can be inserted between memory retrieval and the main LLM loop to perform structured chunk–update–recall–summary reasoning over candidate memories. Heavyweight per-skill dependencies are declared as optional extras in `pyproject.toml` (groups `spatial-domains`, `spatial-histology`, `spatial-wsi`, `memory`, `interactive`, `tui`, `research`) and a runtime `dependency_manager` maps each scientific package to its domain tier, so users install only the dependencies their skills actually need.

### 4.2 Memory mechanism

SpatialClaw’s defining feature is its persistent, structured memory subsystem. Unlike retrieval-augmented agents that maintain only an ephemeral conversation buffer or a flat vector store [15, 16], SpatialClaw stores all dataset metadata, analysis lineage, biological insights, and user preferences in a versioned graph that persists across sessions and can be queried, audited, and rolled back. The implementation lives entirely under `spatialclaw/memory/` and is exposed as both a Python API (for the agent) and a FastAPI REST service (for inspection and external clients).

#### 4.2.1 Graph data model

The memory subsystem is a directed acyclic graph stored in SQLite by default and PostgreSQL when configured for multi-user deployments, with both backends accessed exclusively through asynchronous SQLAlchemy. The schema (`spatialclaw.memory.models`, Table 2) is built from four orthogonal entities. A *Node* (`nodes` table) is a version-independent identifier (UUID) representing a conceptual entity—a dataset, an analysis run, an insight, or a preference—and carries no content itself. A *Memory* (`memories` table) is a single content version of a node, stored as text together with an `is_retired` flag and an optional `superseded_by_id` pointer to its successor; updating content never overwrites a previous version, instead appending a new row that supersedes the old. An *Edge* (`edges` table) is a directed parent → child relationship between two nodes, carrying a display `name`, an integer `priority`, and an optional `disclosure` field that doubles as a typed payload (e.g., a similarity score for `task_similar` edges, see §4.2.5); the (`parent_uuid`, `child_uuid`) pair is unique, so the structural relationship is canonical even when multiple URI aliases reference it. A *Path* (`paths` table) is a materialized routing cache keyed by (`domain`, `path_string`) → `edge_id`; the source of truth for tree structure is the edges table, and paths are a routing convenience.

**Table 1:** Skill metadata schema as declared in SKILL.md frontmatter and the registry entries of spatialclaw.core.registry.

| Field | Type | Required | Description |
| --- | --- | --- | --- |
| **name** | string | yes | Unique skill identifier (kebab-case) |
| **description** | string | yes | Natural-language summary used by the LLM router |
| **version** | semver | yes | Skill version for reproducibility |
| **domain** | enum | yes | spatial / orchestrator |
| **tags** | list[str] | yes | Searchable keywords |
| **trigger_keywords** | list[str] | no | Keywords for the keyword-based router |
| **allowed_extra_flags** | set[str] | no | Whitelisted CLI flags accepted from the LLM |
| **llm_argument_schema** | object | no | Per-skill argument coercion rules (aliases, allowed values, numeric bounds) |
| **requires_preprocessed** | bool | no | Whether input must be a preprocessed Ann-Data |
| **saves_h5ad** | bool | no | Whether the skill produces an updated Ann-Data |

**Table 2:**
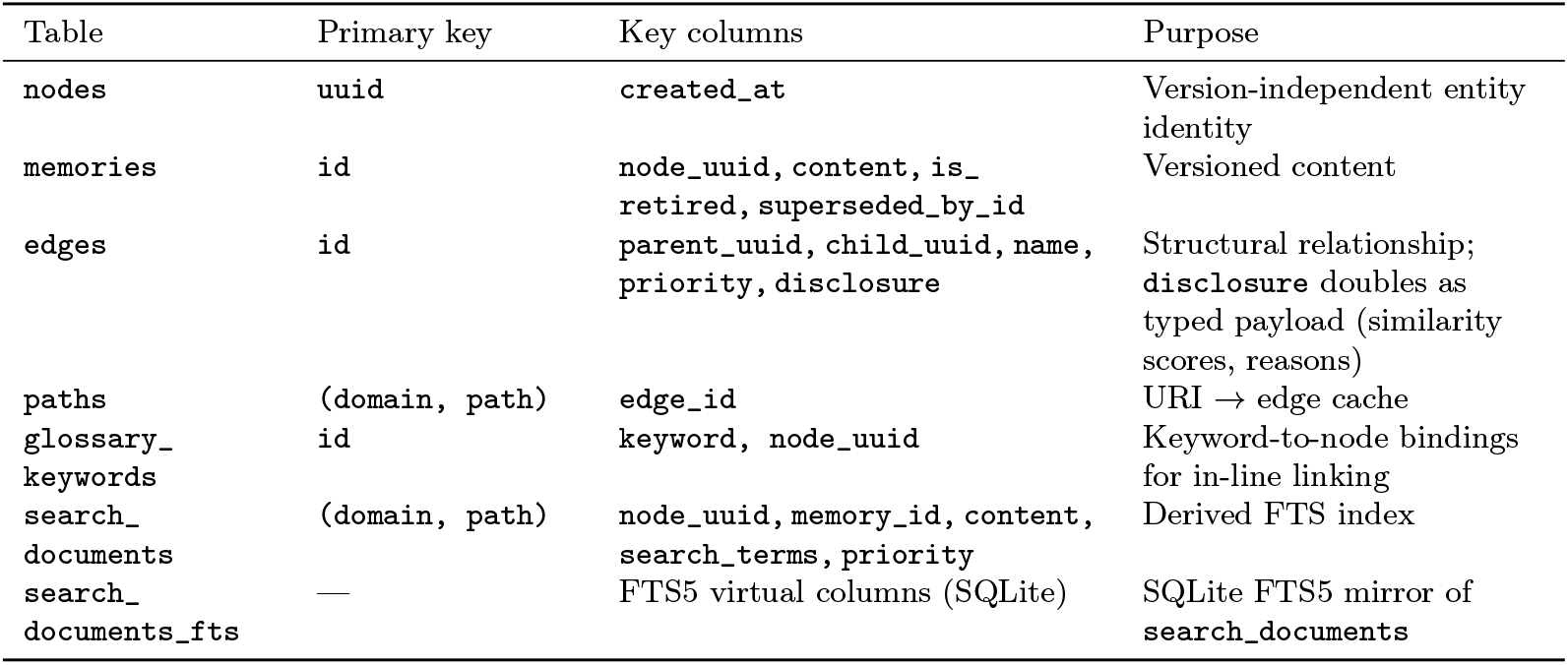
Memory graph schema. All seven tables are defined in spatialclaw.memory.models; the FTS5 virtual table is created at startup by DatabaseManager._create_fts_table when the SQLite build supports it.

| Table | Primary key | Key columns | Purpose |
| --- | --- | --- | --- |
| `nodes` | `uuid` | `created_at` | Version-independent entity identity |
| `memories` | `id` | `node_uuid, content, is_retired, superseded_by_id` | Versioned content |
| `edges` | `id` | `parent_uuid, child_uuid, name, priority, disclosure` | Structural relationship; `disclosure` doubles as typed payload (similarity scores, reasons) |
| `paths` | `(domain, path)` | `edge_id` | URI → edge cache |
| `glossary_keywords` | `id` | `keyword, node_uuid` | Keyword-to-node bindings for in-line linking |
| `search_documents` | `(domain, path)` | `node_uuid, memory_id, content, search_terms, priority` | Derived FTS index |
| `search_documents_fts` | — | FTS5 virtual columns (SQLite) | SQLite FTS5 mirror of `search_documents` |

Memories are addressed externally by URIs of the form `<domain>://<path>`, where the domain partitions the namespace across ten values—`core`, `project`, `dataset`, `analysis`, `preference`, `insight`, `notes`, `session`, `episodic`, `semantic`—and the path encodes hierarchy via slash-separated segments. The same node can be referenced by multiple paths, enabling lightweight aliasing without exposing UUIDs to the LLM. A sentinel root node with the fixed UUID `00000000-0000-0000-0000-000000000000` anchors the graph and ensures uniqueness constraints behave consistently across SQLite and PostgreSQL (which differ in NULL-comparison semantics).

This four-entity separation isolates four concerns that real agentic systems repeatedly conflate [23, 24]: *what* an entity is (Node), *what it currently says* (Memory), *how it relates structurally* (Edge), and *how it is addressed by callers* (Path). Updating content, restructuring the hierarchy, and adding aliases each touch only one of the four tables, simplifying both reasoning and rollback.

#### 4.2.2 Episodic and semantic layers

Long-running agent design must decide which experiences to retain as transient session traces and which to elevate to durable, transferable knowledge [21]. SpatialClaw maintains two parallel domain spaces in the graph: an *episodic* layer (`episodic://…`) for raw session events, and a *semantic* layer (`semantic://…`) for consolidated knowledge. The same memory object can simultaneously exist in both layers, each version stamped with layer-specific metadata (`importance`, `confidence_score`, `evidence_uri`, `promoted_at`, and human-readable `promotion_reasons`). At retrieval time, callers may scope queries to a single layer or to both via the `normalize_memory_layer` helper.

The platform exposes five typed memories (Pydantic models in `spatialclaw.memory.store`): `DatasetMemory` (file path, platform, *n*_obs_, *n*_vars_, preprocessing state), `AnalysisMemory` (skill, method, parameters, output path, status, duration, task summary, key steps, failure reason, dataset lineage), `PreferenceMemory` (domain, key, scalar value, strict flag), `InsightMemory` (entity type/id, biological label, evidence, AI-predicted vs. user-confirmed flag, learned rule, score, supporting/refuting task URIs), and `ProjectContextMemory` (project goal, species, tissue type, disease model). Whether an episodic memory is also written to the semantic layer is decided at write time by a deterministic, type-aware policy `evaluate_promotion` (`spatialclaw.memory.promotion`). Each candidate memory is scored on two clamped-to-[0, 1] dimensions—*importance* (long-term reuse value) and *confidence* (factual reliability). Both scores accumulate from type-specific signals and are returned together with natural-language reasons (Table 3). The rules favour precision over recall, since polluting long-term memory is more harmful than leaving a useful fact in episodic-only storage. User preferences and project context are inherently semantic and bypass the episodic layer entirely. Analysis records are promoted only if `status == “completed”` and they carry either an output path or a dataset-lineage pointer. Insight records require either user confirmation or an attached evidence string. Every promotion decision, including the reasons that triggered it, is preserved in the memory itself for downstream auditing.

**Table 3:** Promotion rules by memory type, as implemented in evaluate_promotion (spatialclaw. memory.promotion). importance and confidence are accumulated from type-specific signals and clamped to [0, 1].

| Memory type | Default layer | Promotion trigger |
| --- | --- | --- |
| **preference** | semantic only | Always |
| **project_context** | semantic only | Always |
| **dataset** | episodic | $\text{confidence} \geq 0.75 \wedge \text{importance} \geq 0.55$ |
| **analysis** | episodic | $\text{status} == \text{completed} \wedge (\text{output\_path} \vee \text{source\_dataset\_id})$ |
| **insight** | episodic | $\text{confidence} == \text{user\_confirmed} \vee \text{explicit evidence}$ |
| **unknown** | episodic only | Never |

#### 4.2.3 Storage path: remember

The agent writes to memory exclusively through the high-level `MemoryClient` (`spatialclaw.memory. memory_client`), which exposes five verbs—`remember`, `recall`, `forget`, `search`, `boot`—over the URI model above. A single `remember(uri, content, priority, disclosure)` call (i) parses the URI into `(domain, path)`, (ii) consults the `paths` table to decide whether the URI already resolves to an active node, and either (iii_a_) appends a new `Memory` version on the existing node and updates edge metadata, retiring the previous version via `is_retired = True` and `superseded_by_id`, or (iii_b_) creates a new node, materializes the parent path chain up to the sentinel root, inserts a new edge, and registers a new path. Across both branches the `GraphService` layer (`spatialclaw.memory.graph`) emits before/after row pairs to a `ChangeCollector`, which the client forwards to the `ChangesetStore` (§4.2.6). The typed-memory facade `SpatialMemoryStore.save_memory` sits on top of this primitive: it computes the promotion decision, stamps layer-specific metadata, and issues one `remember` call per eligible layer, so a single semantically equivalent memory may be written to both `episodic://` and `semantic://` URIs within one transaction.

#### 4.2.4 Retrieval path: recall **and** search

Retrieval is offered along two complementary axes. *Direct recall* resolves a known URI to its current active memory in a single indexed lookup against the `paths` table; this is the fast path used when the agent already holds a reference to a specific entity (e.g., a session ID or a known dataset). *Full-text search* is served by a derived `search_documents` table (`spatialclaw.memory.search`) that mirrors every reachable `(node, path)` pair together with content, disclosure, and an auxiliary `search_terms` field. The latter is populated by a tokenizer (`spatialclaw.memory.search_terms`) that normalizes URI separators, segments CJK runs through `jieba` (with bioinformatics terminology auto-registered as custom words), and concatenates path tokens, content tokens, and glossary keywords. Two back-ends are supported transparently: SQLite FTS5 with a tuned BM25 weighting that emphasizes path and URI matches over content (`bm25(0.0, 2.5, 0.0, 2.0, 1.0, 1.0, 0.75)`), and PostgreSQL `tsvector`/`ts_rank_cd` for production deployments. A failed FTS5 query (e.g., due to an older SQLite build) falls back to a LIKE-based scan over the same documents. Mutations trigger `refresh_search_ documents_for_node`, which deletes and re-materializes only the affected node’s rows, avoiding global re-indexing.

To bridge unstructured content and the underlying graph, each node may be annotated with one or more glossary keywords (`glossary_keywords` table). New memory content is scanned by an Aho–Corasick automaton (`spatialclaw.memory.glossary`) that is lazily rebuilt against a fingerprint of the keyword table—row count, maximum id, maximum `created_at`—and returns every keyword occurrence with its bound node, allowing front-end clients to render in-line links from free text to structured graph entities.

A higher-level `SpatialMemoryStore.load_context(session_id)` call assembles the LLM prompt’s memory block by composing layer- and type-scoped queries: at most one semantic project context, one episodic dataset, three recent analyses (drawn from both layers), five semantic preferences, and three semantic insights. During workflow continuation, a `filter_context_preferences_for_ continuation` helper strips workflow-state preferences from the assembled block to prevent stale defaults from overriding the most recent completed analysis.

#### 4.2.5 Insight extraction and graph extension

Beyond passively storing experiences, SpatialClaw includes an `InsightExtractor` (`spatialclaw. memory.insight_extractor`) that periodically distills reusable analytical rules from accumulated `AnalysisMemory` records. This self-evolving extraction mechanism draws on recent work in graph-based experience augmentation [26] and reward-driven memory consolidation [27]. After every five completed tasks (`START_THRESHOLD = ROUNDS_PER_FINETUNE = 5`), the extractor enters a finetune phase: it samples up to `INSIGHTS_POINT_NUM = 5` anchors from successful and failed analyses, retrieves currently active insights whose `positive_task_uris` overlap with the sampled task summaries above a threshold of half the sample size, and prompts the LLM with two structured prompts—a comparison prompt pairing a successful and a failed trajectory, and a pure-success prompt—asking it to update the rule set with one of four operations: Add, Edit, Remove, or Agree. Each operation is parsed back into the graph as either a new `InsightMemory` node or an in-place update.

The graph is extended at this point with two additional edge types that reuse the standard `edges` schema. When a new analysis is recorded, `remember_analysis_with_similarity` computes Jaccard similarity between the new task’s `task_main` summary and those of all prior analyses in the same domain, creating undirected `task_similar` edges (with the similarity score stored verbatim in the edge’s `disclosure` field) for pairs above a configurable threshold (default 0.7). At retrieval time, `get_task_neighbors` performs a bounded *k*-hop breadth-first traversal from a query node along these similarity edges, optionally filtered by a minimum similarity. A second edge type, `derived_from`, is created from each insight to the analyses that supported it, preserving full provenance. Combined with the FTS index, a single retrieval call can therefore return successful exemplars, failure cases, and the rules learned from both—a richer context than either pure vector retrieval or pure FTS could provide.

Insights carry a numeric `score` initialized at 2.0, incremented when the LLM Agrees with an existing rule and decremented when it Removes. A `backward(reward)` API allows downstream task outcomes to reinforce or penalize the insights consulted during planning; insights whose score decays to zero are pruned. A periodic `merge_insights` operation clusters analyses by `cluster_id` and asks the LLM to deduplicate the rules associated with each cluster, capping the rule set at `MAX_RULE_NUM = 10` per cluster and preventing rule-set bloat over long-running deployments.

#### 4.2.6 Auditability and rollback

Because the agent autonomously mutates memory on the user’s behalf, every write must remain inspectable and reversible. A `ChangesetStore` (`spatialclaw.memory.snapshot`) accumulates row-level before/after pairs across all mutations into a single JSON-backed pool persisted at ∼`/.config/spatialclaw/memory_snapshots/changeset.json` (overridable via `SPATIALCLAW_ SNAPSHOT_DIR`). The store applies overwrite semantics: the first time a primary key is touched within an audit window, both *before* (pre-AI) and *after* (post-AI) states are captured; subsequent touches update *after* only, freezing the original *before* for review. Net-zero changes (*before* == *after* ) and create-then-delete no-ops are filtered automatically by a garbage-collection pass that also removes orphaned dependents. The JSON pool itself is written via a temporary file plus `os.replace` to guarantee filesystem-level atomicity even on interrupted writes.

A FastAPI server (`spatialclaw.memory.server`) exposes the graph and the changeset trail through three router groups, all optionally protected by a bearer token configured via `SPATIALCLAW_ MEMORY_API_TOKEN`. `/api/browse` provides read/write access to nodes, paths, children, recent memories, full-text search, and glossary entries. `/api/review` surfaces the changeset store—change list, change count, per-key diff, and rollback—so that a human reviewer can inspect every AI-induced mutation and selectively revert it. `/api/maintenance` lists retired memory versions, supports their permanent deletion, and exposes a search-index-rebuild endpoint for catastrophic recovery. Rollback is symmetric: reverted content updates restore the previous `Memory` version through `rollback_to_memory`; reverted path creations are handled by `remove_path`; reverted path deletions are reconstructed via `restore_path`. All graph writes are atomic at the SQLAlchemy session level, so a failed mutation cannot leave the changeset and the database in inconsistent states.

### 4.3 Reproducibility

SpatialClaw enforces reproducibility at three levels: the environment, the individual analysis run, and the multi-step session.

#### Environment

The package targets Python 3.10–3.12 and is distributed through `pyproject.toml` with a small core dependency set (Scanpy ≥ 1.9, AnnData ≥ 0.11, Squidpy ≥ 1.2, MuData ≥ 0.2, plus standard scientific Python libraries) and seven optional extras—`spatial`, `spatial-domains`, `spatial-histology`, `spatial-wsi`, `memory`, `interactive`, `tui`, and `research`—that pull in only the heavyweight packages a given workflow actually requires (e.g., PyTorch and PyTorch Geometric for graph-neural-network domain identification methods, `aiosqlite` and FastAPI for the memory sub-system, `prompt-toolkit` and `textual` for the interactive layer, LangChain and DeepAgents for the research pipeline). A runtime `dependency_manager` (`spatialclaw.core.dependency_manager`) maps each scientific package to its domain tier and is used by skills to surface clear, actionable error messages when an extra is missing rather than failing deep inside an import chain.

#### Per-skill reproducibility bundle

Every skill writes to its output directory a fixed contract described in §4.1: a Markdown narrative `report.md`, a machine-readable `result.json` envelope, an updated `processed.h5ad` (when applicable), plots under `figures/`, tables under `tables/`, and a `reproducibility/` sub-directory with the exact command that was run, an environment specification (`environment.yml`), and a SHA-256 checksum file (`checksums.sha256`) covering all inputs. The `result.json` envelope itself records the skill’s semver version, completion timestamp, and input check-sum, so a later run can be unambiguously matched against a prior result. Reports include the standard SpatialClaw disclaimer that the autonomous ecosystem is a research and educational tool and not a clinical device.

#### Per-session reproducibility

The `AnalysisSession` object (`spatialclaw.common.session`) persists session metadata (input file path, dataset SHA-256 checksum, species, data type, domain, creation timestamp) together with every skill’s accumulated result envelope to a single JSON file. Saving and loading a session is symmetric, so a multi-step workflow can be paused, resumed in a different process, or shared with a collaborator while preserving the exact dataset binding, processing-state flags, and historical skill outputs. The persistent memory subsystem of §4.2 adds a complementary, agent-level reproducibility guarantee: every analysis is recorded as an `AnalysisMemory` with skill, method, parameters, output path, status, duration, and the LLM-readable `task_main` and `key_steps` summaries, so the autonomous ecosystem retains an auditable trace even when individual session JSON files are lost.

#### Privacy and data boundaries

By design, the memory subsystem stores metadata, parameters, path summaries, analysis state, report excerpts, and biological conclusions, but never the raw expression matrices themselves. `DatasetMemory` explicitly forbids absolute paths (enforced by a Pydantic validator), and the bot’s auto-capture path reduces external paths to their basename before persistence to avoid leaking server-side directory structures. `PreferenceMemory` is constrained by a separate validator (`preference_policy`) that rejects paths, URLs, multi-line content, and structured payloads, ensuring that workflow facts are recorded as `dataset`, `analysis`, or `insight` memories rather than polluting user preferences.

## 5 Acknowledgements

This work is supported by the National Natural Science Foundation of China [62433016].

## 6 Code availability

SpatialClaw is open-source under the MIT license at https://github.com/ShangBioLab/SpatialClaw.

## Notes

https://github.com/ShangBioLab/SpatialClaw

### Competing Interest Statement

The authors have declared no competing interest.

